# 3D bioprinting of low-viscosity phase-separated food-grade bioinks by *in situ* self-assembly

**DOI:** 10.1101/2024.07.06.602353

**Authors:** Sara M. Oliveira, Gabriel DeSantis, Lorenzo M. Pastrana

## Abstract

There is a notable gap in the scientific understanding of the cellular role in cultured cell-based foods. Unravelling the effects of the interactions between ingredient micro/nanostructure and cells and their significance on nutrition and texture is of great importance. In addition, bioprinting methods face notable limitations in animal-free formulations and scale. Herein, we introduce a proof-of-concept bioprinting method based on the *in situ* integration of self-assembling events, allowing printing without supporting baths. Our approach enabled a food-grade 3D bioprinted model with 8.5 mm height and a hardness of 284 mN, supporting the early differentiation of myoblasts producing embryonic myosin heavy chain, after 7 days of differentiation. Cellular protein content increased up to 18-fold per initial cell without changes in construct texture. The method provides a novel concept to produce robust, cell-dense platforms for further research on food-grade bioprinted foods.

## 1 Introduction

Bioprinting living cells holds great promise in enhancing and personalizing cell-based food products (Guo et al., 2023; Ianovici et al., 2022; Ma & Zhang, 2022). It enables the incorporation of higher cell concentrations than scaffold-based approaches and customization of the construct’s nutritional and organoleptic properties. Despite this, several limitations remain, such as printing mechanisms and food-grade formulations that enable high cell viability and efficient protein production. New solutions should also be designed to increase bioprinting mass flow, which is mainly defined by the printing speed and nozzle diameter – **Fig. 1a**. Furthermore, there are not yet scalable and validated solutions for food-grade bioprinting. **Table 1** summarizes the current scientific publications on cultivated meat/fish applications. The majority of these still use animal-derived ingredients and/or do not provide data on the textural or nutritional qualities of the constructs

**Table 1.**
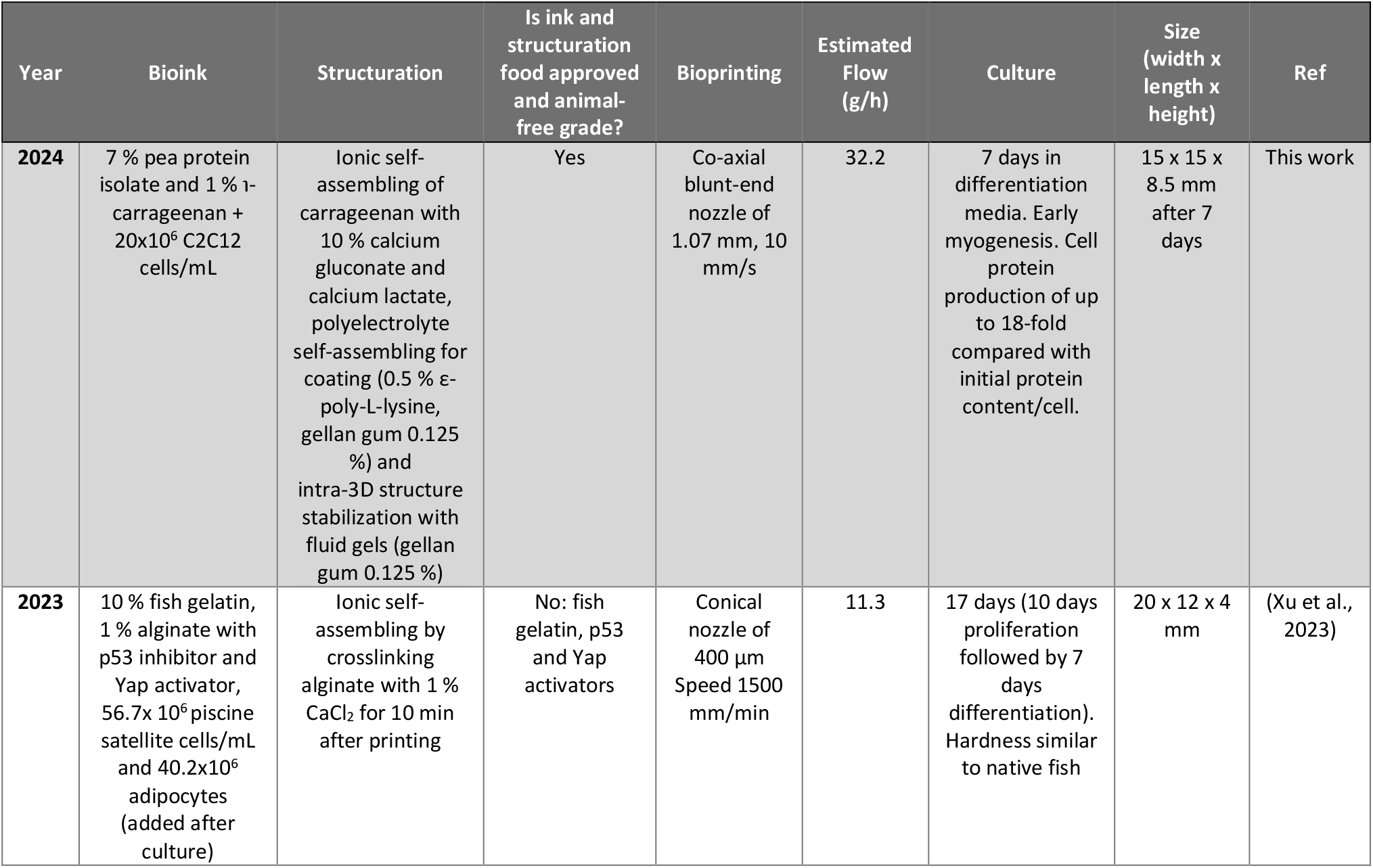
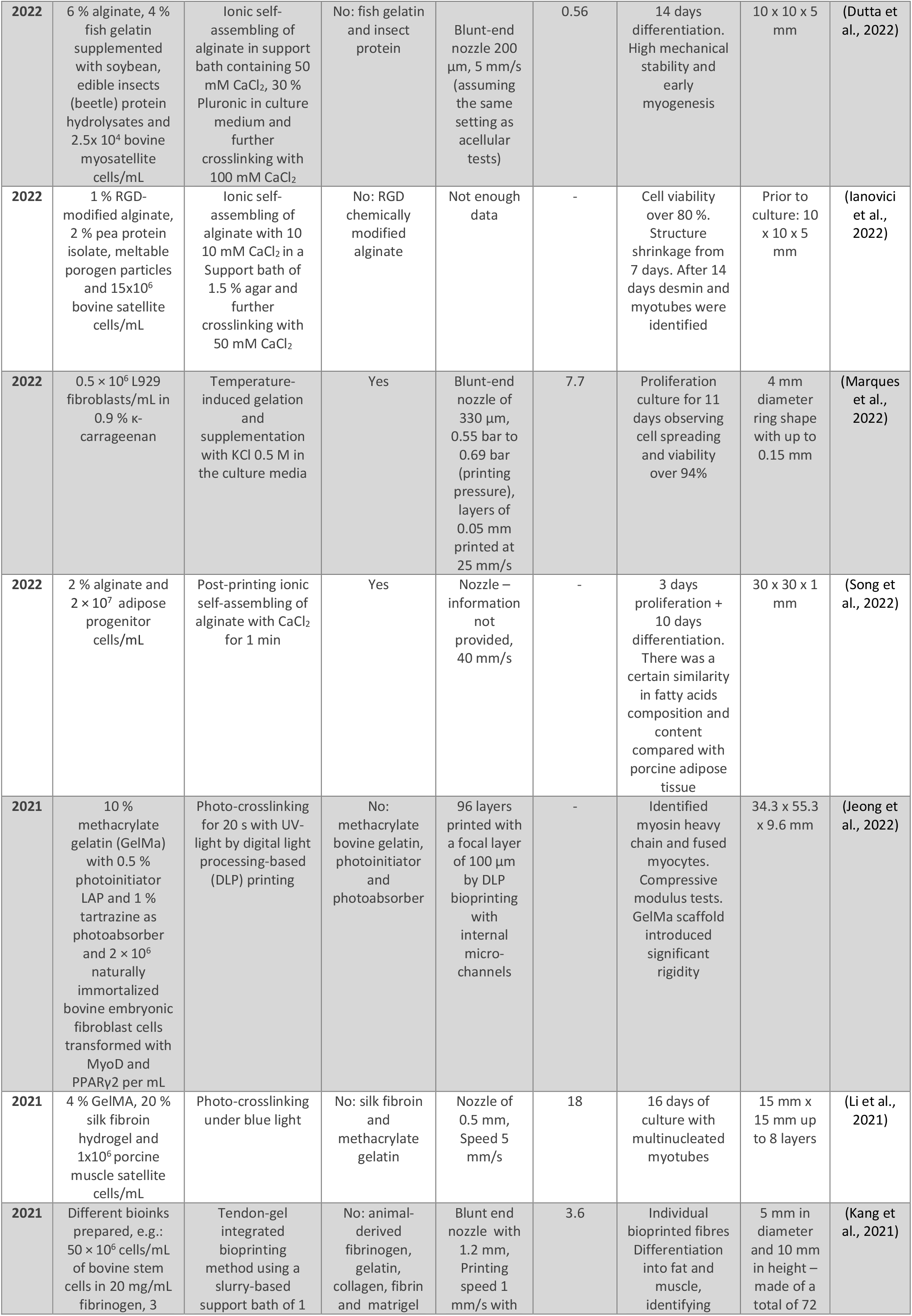
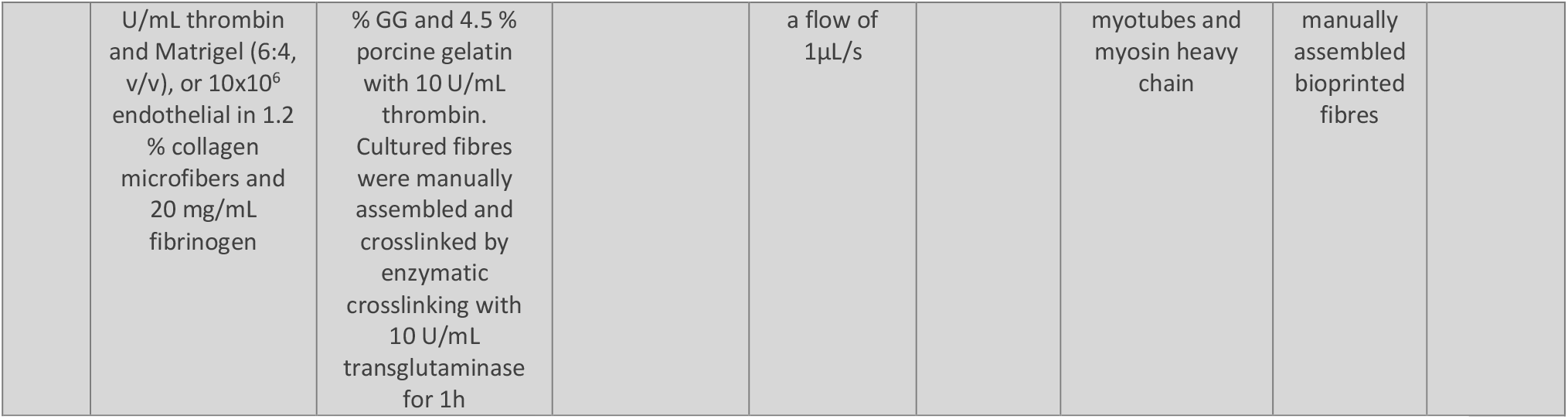
Comparative analysis of bioprinted cultivated meat studies reported in the literature. The mass flow was estimated assuming a circular cross-section area multiplied by the printing speed and the bioink density similar to water density (998 kg/m^3^).

**Fig. 1.**
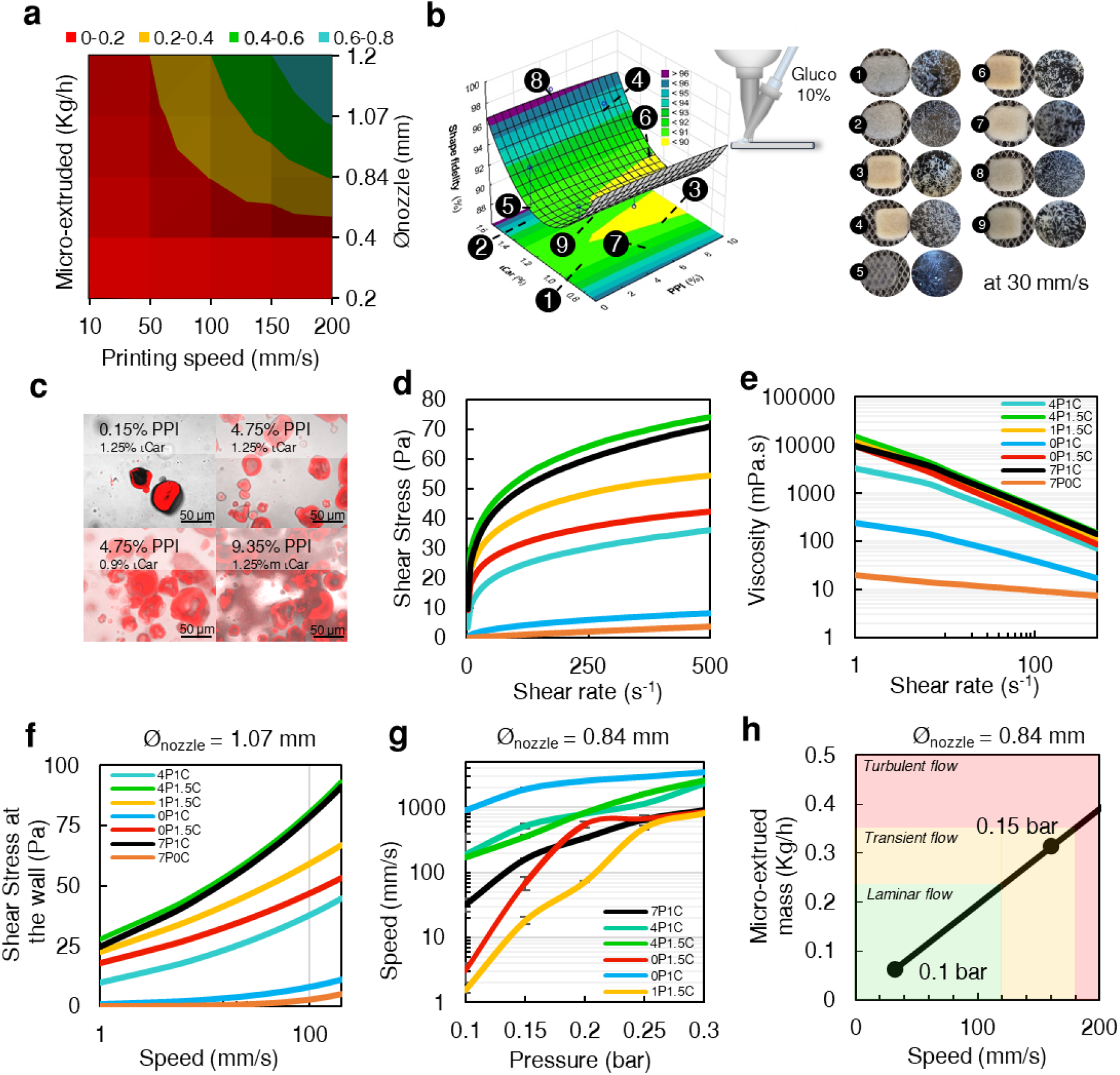
Printing, structuration and characterisation of the food-grade inks. (a) Estimated maximum micro-extruded mass for different printing speeds and nozzle diameters, assuming a 998 kg/m^3^ density and a circular cross-section. (b) Surface response illustrating the impact of PPI and ℩Car concentration on the shape fidelity of the 3D structures printed with Gluco 10 % deposited drop-wise and printed at 30 mm/s with a 0.6 mm nozzle with 100 % infill. Photographs of the printed samples alongside the disrupted samples confirm the formation of microfibre-like structures. (c) Confocal images of the pea microstructures stained with Rhodamine B. (d,e) Flow curves depicting the effect of shear rate on shear stress and viscosity, respectively (1-500 s^-1^). (f) Extruded mass at different pressures plotted against estimated printed speeds. (g) Estimated shear stress at the wall varying printing speeds. (h) Relationship between micro-extrusion pressure and mass, flow regime and the estimated printing speed for 7P1C ink.

The lack of scalable and validated solutions is also likely due to the variety of steps required in establishing a bioprinting process, such as: 1) cell expansion, 2) bioinks fabrication, 3) bioprinting, 4) culture of the bioprinted products, 5) post-processing, and 6) packaging. Significant complexity in the process originates from several physico-chemical factors through the process stages that influence cell behaviour and can also jeopardize cell viability and alter cell function (Zhao et al., 2015). Furthermore, controlling and optimizing cellular activity (e.g., adhesion, death, spreading, fusion, differentiation, protein or matrix production) in these different process stages is crucial to reducing production time and improve product quality. In addition, controlling shear stress is a fundamental factor for cell integrity and these other downstream cellular processes but generally runs counter to the desired criteria of high print speeds (Blaeser et al., 2016). While highly viscous or highly crosslinked bioinks exhibit excellent representation of the 3D sliced model (i.e., high printability), they will induce cell trapping, reduce mobility and might limit intracellular protein production. Thus, low-viscosity, thixotropic and shear-thinning bioinks have emerged as popular ink types to reducing the shear stress required to print robust, high-fidelity constructs (Liu et al., 2017). However, achieving 3D bioprinted structures with distinguishable fibres, shape stability, and no delamination during the culture period is extremely challenging.

The most common method to print low-viscosity inks is based on the Freeform Reversible Embedding of Suspended Hydrogels (FRESH) method, initially developed by Adam Feinberg *et al*. (Hinton et al., 2015). It consists of printing in a slurry-based support bath, usually consisting of densely fractured gels made from gelatin, agarose, alginate, or high-viscosity solutions. They can be reversibly liquefied and combined with curing agents, enabling sample crosslinking and removal from the slurry. Notwithstanding, using long needles is a scale-out limiting factor in speed and maximum printable height. At high speeds, the needle bends, creating turbidity in the printing media (Mirdamadi et al., 2020).

We consider that new structuration methods capable of printing soft 3D structures without supporting baths could reduce scalability limitations. Compared with FRESH-based methods, printing resolution might be reduced; however, production capacity is a priority for cell-based food.

Bioprinting soft structures could enhance myoblast mobility and protein production, which could be particularly relevant for increasing culture efficiency, i.e., the protein production capacity per cell. This is especially significant in meat, where nearly half of the protein is located intracellularly (Lawrie & Ledward, 2014; Voutila, 2009).

Prior work suggests that the *in situ* integration of technologies could increase the multiscale control over 3D constructs and improve cell behaviour (Oliveira et al., 2015). With this concept in mind, the main objectives of this work were: i) to formulate and characterize new low-viscosity food-grade bioinks; ii) to develop a novel *in situ* structuration mechanism for the low-viscosity inks, with effective nutrient delivery and scalability in mind, via the fabrication of bioprinted structures with open pores, and not using support baths; iii) to validate the capacity of the new inks to support cell activity; iv) to produce a stable soft bioprinted model sufficiently high for texture profile analysis; v) to preliminarily assess the impact of cells and their differentiation potential regarding protein content and texture.

## 2 Materials and Methods

### 2.1 Materials

Several protein isolates/concentrates were obtained from Bulk: whey protein isolate (WPI, 97g/100g), pea protein isolate (PPI, 80g/100g protein) and Brown rice protein (RP, 80g/100g protein), soy protein isolate (SPI, protein 85g/100g protein). Gellan Gum (Gellan), τ-carrageenan (Iota), κ-carrageenan (Kappa) and calcium gluconate & calcium lactate (Gluco) were purchased from Texturas Ferran Adriàn.

### 2.2 Formulation of inks

Stock dispersions of 2.5% polysaccharide (w/v) and 20% protein (w/v) were prepared by dispersing the solids in deionized water. All dispersions were used within 24 hours. The protein and polysaccharides were mixed in different proportions. Immediately after mixing, the dispersions were homogenized by vortexing at 3000 rpm for 1 min. The samples were heated at 86°C for 1 hour in a thermomixer to induce gelation and phase separation. The samples were cooled in ice and vortexed until the gelation initiation was observed. The samples were stored at 4°C. The protein concentrations tested were 1, 5, 10 and 20%, and the polysaccharides concentrations were 0.5, 1, 1.5 and 2%. Low protein concentration (1%), and high protein concentration (10%) were combined with 1 or 1.5% polysaccharides-Fig. S2.

### 2.3 Printing tests and shape fidelity

For the starting qualitative printability tests – Fig. S1 and Fig. S2, a cylinder (20 mm diameter with 5 mm height) was designed with AutoCAD (v. 2013) and exported as an STL file. The G-code was generated using simplify3D (v 4.1.2). The model was sliced for a nozzle diameter and extrusion width of 1.2 mm (tapered tip), layer height of 0.84 mm, multiplier of 0.8 and a printing speed of 30 mm/s. For the quantitative analysis made through Desing of Experiments (DoE) and the printing tests displayed in Fig. 1, a prism (2 x 2 cm with a height of 4 mm) was sliced for a nozzle diameter of 0.6 mm (tapered tip), extrusion width of 0.6 mm, 0.6 mm filament width, 0.8 flow multiplier, 0.287 mm layer height and printing speed of 30 mm/s. The G-codes were uploaded to a Focus 3D Food Printer (byFlow, the Netherlands).

The model used for the DoE was the central composite model, defined by:

, whehere Z, printability and X_1_, X_2_, are the coded values of the independent variables (*i*.*e*. protein concentration, polysaccharide concentration). X_i_, X_i_^2^ and, X_i-1_·X_i_ identify the interactions between the variables; β_i_, β_ii_, β_i-1,i_ are the coefficients of the mentioned variables and the interactions; and β_0_ is the independent term. The runs and the natural values of the independent and dependent variables of the central composite experimental design are presented in **Table S1**. The protein concentration was varied between 0.15 to 9.35 %, and the polysaccharide between 0.9 and 2.5 %.

where Z, printability and X_1_, X_2_, are the coded values of the independent variables (*i*.*e*. protein concentration, polysaccharide concentration). X_i_, X_i_^2^ and, X_i-1_·X_i_ identify the interactions between the variables; β_i_, β_ii_, β_i-1,i_ are the coefficients of the mentioned variables and the interactions; and β_0_ is the independent term. The runs and the natural values of the independent and dependent variables of the central composite experimental design are presented in Table S1. The printing was performed using a droplet-based and custom-made nozzle with a 0.6 mm nozzle (to micro-extrude the ink) and a 1.2 mm nozzle to dispense the ionic crosslinked solutions selected based on preliminary tests (10 % Gluco). The flow rate of Gluco was controlled using a syringe pump with a syringe with a diameter of 26.59 mm (50 mL syringe) and a feeding flow rate of 2.1 mL/min. The actual length, width and height of the printed samples were measured using a calliper and used to calculate the shape fidelity (*i*.*e*., the Z response) based on the following equation (Dutta et al., 2022):

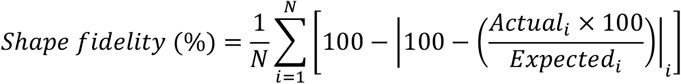

The samples were photographed from different angles after printing using a 12-megapixel camera with a resolution of 4619 × 3654 pixels of a Samsung Galaxy Note 10+ (f/1.5–2.4, 1–1.4 μm pixels, 34.7/4.3 mm focal length, 495 ppi with dual-pixel autofocus and optical image stabilization) and the holders displayed in Fig. S1 and Fig. S3.

Analysis of variance (ANOVA) was used to determine the significance and adequacy of the model using Statistica 10 (Tibco). Each condition was assessed in triplicate samples and three central points.

### 2.4 Confocal microscopy

The samples were fixed with a 4 % formalin solution for 15 min at room temperature. The fixative was removed, and the samples were rinsed with PBS thrice. The samples were incubated with 0.2 % (w/v) rhodamine B solution in PBS for 25 min in the dark. And, then were extensively rinsed with PBS before analysis. The samples were observed by <u>confocal microscopy</u> (Zeiss LSM 780) and excited at the wavelengths of 488 and 561 nm. Emission was set at 570–620 nm, including T-PMT, and the observation was carried out with 40 × and 63 x oil Zeiss objectives.

### 2.5 Rheology

A stress-controlled Rheometer MCR 302e (Anton Paar) was used for the rheological analysis. The viscosity recovery capacity and thixotropic behaviour were measured by performing the three-interval thixotropy test using a <u>stainless steel</u> flat plate geometry (50 mm, 1000 μm gap) at 24 °C. The three intervals defined were: 1) shear rate of 0.25 s^-1^ for 20 s, 2) shear rate of 1000 s^-1^ for 1 s, and 3) shear rate of 0.25 s^-1^ up 500s. All the samples were pre-sheared at 0.25 s^-1^ for 20s. Flow curves were performed using a <u>stainless steel</u> flat plate geometry (50 mm, 1000 μm gap) at 24 °C. Flow curves were obtained using an up-down-up step-wise program with the shear rate ranging from 1 to 500 s^−1^. The inks were pre-sheared (increasing, decreasing and increasing shear rate), and the third curve was used for the analysis. The flow curves were fitted to the Herschel-Bulkley model, which is given by:

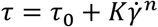, where *τ*_0_ is the yield stress at zero shear rate (Pa.s), *K* is the consistency coefficient (Pa.s^n^), and *n* is the flow behaviour index (dimensionless). The theoretical printing flow was calculated with the equation

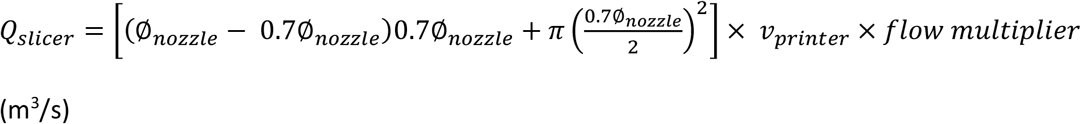

where ∅_nozzle_ is the nozzle diameter (m); h is the layer thickness = 0.7*∅_nozzle_ (m); *v*_*printer*_ is the printing speed (m/s); w is the filament width = ∅_nozzle_ (m); and *flow multiplier* was consider 1. The corrected shear stress at nozzle wall, was calculated by combining the Herschel-Bulkley model and the Rabinowitch correction for non-Newtonian fluids, resulting:

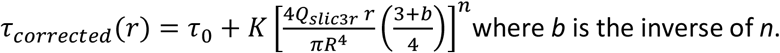

### 2.6 Micro-extrusion at controlled pressure

A pressure-controlled pneumatic system (DC-100 Pressure Controller, Fisnar) equipped with a tapered nozzle of 0.84 mm diameter and a 50 mL barrel was used to quantify the extruded ink. Pressures ranging from 0.1 to 0.3 bar were applied for 3 s and the extruded mass was weighted.

Subsequently, the mass was converted to the theoretical printing speed, considering the circular extrusion area of the nozzle.

The densities of the inks were calculated by weighting 1 ml of the inks:

| PPI (%) | Car (%) | Density (g/mL) |
| --- | --- | --- |
| 4 | 1 | 0.97530 |
| 4 | 1.5 | 0.96913 |
| 1 | 1.5 | 0.99990 |
| 0 | 1 | 0.97280 |
| 0 | 1.5 | 1.00150 |
| 7 | 1 | 0.98063 |

### 2.7 Reynolds number

The Reynolds number is given by:

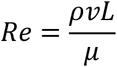

Here, *ρ* is the density of the fluid (Kg/m^3^), *v* is the ink speed (m/s, considered equal to *v*_*printer*_), *μ*is the ink dynamic viscosity (Pa.s), *L* is the characteristic length or the hydraulic diameter of the pipe, considered as the nozzle diameter, ∅_nozzle_ (m). Flow is considered laminar if *Re*< 2000, transient between 2000-4000 and turbulent above 4000. We have considered that the dynamic viscosity at a given printing speed corresponds to the corrected apparent viscosity at the wall. Based on the Herschel-Bulkley model and the Rabinowith correction for non-Newtonian fluids, the corrected dynamic viscosity is:

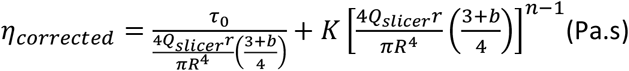

Here, *Q*_*slicer*_is the micro-extruded flow rate (m^3^/s), *R* is the nozzle radius (m), *r* is the point in the transversal profile of the nozzle cross-section (m) – at the wall *r*=*R*, and b is the inverse of the flow index (1/*n*). Combining and simplifying both equations, *Re* can be estimated by:

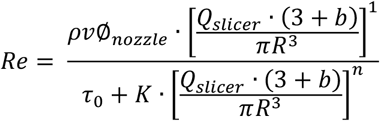

Assuming that the inks are non-compressive fluids, the ink density *ρ* was considered constant.

### 2.8 In situ self-assembling with PLL

Food grade, ultrapure ε-Poly L-Lysine was purchased from Handary (PLL, 2211). Petri dishes were filled with 10% Gluco combined with different concentrations of PLL (5, 15, 25 and 50 mg/mL). All the solutions were prepared in de-ionized water. 20 μL of the ink was dispensed in the crosslinking media. The samples were removed after 1, 3, 5 and 10 min. The particles were disrupted by squeezing with aluminium foil, and the excess liquid was removed with tissue paper. The remaining solid part, which had a whitish colour, not resembling the ink, was weighted.

The effect of PLL concentration in the shape of printed samples with 100 % infill (volume and top area) was calculated following the previous printing settings, except that the external dimensions of the 3D shape were 1×1×1 cm, and the nozzle used was a co-axial with an internal diameter of 1.07 and an external diameter of 1.8 m. The 7P1C ink (50 mL barrel) was structured with Gluco or both Gluco/PLL. The flow rate of the crosslinking solution was controlled using a syringe pump with a syringe with a diameter of 26.59 mm (50 mL syringe) and a feeding flow rate of 2.1 mL/min. The samples were incubated at 37 °C for up to 6 days, and the top surface area change was calculated.

In order to evaluate the effect of In situ self-assembling with PLL/Gluco and GG in a single-wall and porous 3D structure (method 3), a single-wall prism (1.07 width) with a porous structure was designed – Fig.3 d. The G-codes were uploaded to a Focus 3D Food Printer (byFlow, the Netherlands). The extrusion multiplier was set to 0.95, the layer weight was 0.7 mm, the extrusion width was 1.07 mm, and the printing speed was 10 mm/s. The ink was extruded with the previously described 1.07 mm co-axial nozzle. The flow rate of the crosslinking solution (10 % Gluco and 0.5% PLL) was controlled using a syringe pump with a syringe with a diameter of 26.59 mm (50 mL syringe) and a feeding flow rate of 2.1 mL/min. During the printing, the crosslinking solution was pumping, and then after each layer concluded, approximately 200 μL of a fluid gel composed of 0.12 5% GG (heated and cooled at 86 °C for 1 hour) were deposited in the printed layers using a 5 mL plastic Pasteur pipette. After printing, the samples were photographed from different angles using a 12-megapixel camera with a resolution of 4619 × 3654 pixels on a Samsung Galaxy Note 10+ (f/1.5–2.4, 1–1.4 μm pixels, 34.7/4.3 mm focal length, 495 ppi with dual-pixel autofocus and optical image stabilization).

### 2.9 Stability of self-assembled 3D structures

Unrinsed and PBS-rinsed samples were incubated in PBS/0.2 % sodium azide for up to 7 days at 37 °C inside cell strainers. After each time, the PBS and the liquid excess were removed, and the samples were weighed. Samples at different time points were freeze-dried to measure the dry mass. The moisture was calculated in the PBS-rinsed samples using the following equation:

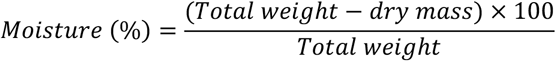

### 2.10 C2C12 proliferation and differentiation

The C2C12 mouse myoblast cell line (ATCC, CRL-1722) was expanded in high glucose, no glutamine, no phenol red and no sodium pyruvate Dulbecco’s Modified Eagle Medium (DMEM, Gibco, 31053028), supplemented with 10 % heat-inactivated Certified Fetal Bovine Serum (FBS, Gibco, 16000044), 200mM GlutaMAX (Gibco, 35050061), 10 mM N-2-hydroxyethylpiperazine-N-2-ethane sulfonic acid (HEPES, Gibco 15630080) and 1 % penicillin-streptomycin (PS, P4333, Sigma-Aldrich), in a humidified atmosphere with 5% CO2 at 37 °C. For expansion, 80-90 % of cell cultures were split into a ratio of 1/3 to 1/6 using Tryple Express Enzyme (Gibco, 12604021) and Dulbecco’s phosphate-buffered saline (DPBS, GIbco no calcium, no magnesium, 14190136). Cell culture was done in Thermo Scientific Nunc™ TripleFlask™ treated Cell Culture Flasks with a 500 cm^2^ growth area.

To induce myogenic differentiation of C2C12, the DMEM supplementation was changed to 2 % Horse serum (HS, GIbco, 16050122), 1 % PS, 10 mM HEPES 1M, 200 mM Glutamax and Insulin-Transferrin-Selenium Liquid Media Supplement (ITS, Sigma-Aldrich, I3146). C2C12 cells were used in passages between 12-26. The culture medium was refreshed every 2-3 days.

### 2.11 C2C12 cytocompatibility in PLL coated PPI-Car

The bioinks were formulated as follows: 10×10^6^ C2C12 cells, 10 μL of DMEM, 25 μL of D-mannose (64.8 % D-mannose in PBS) and 455 μL of ink, for a total of 0.5 mL. The bioinks were dispensed in crosslinking media composed of 1 %PLL and 10 % Gluco for 5 minutes and then washed with DMEM. Cultured media was refreshed each 2-3 days.

### 2.12 Bioprinting of C2C12

The bioprinting was performed in a sterile flow chamber, following method 3 described above in section 2.8. All surfaces, materials, and reagents were previously disinfected (UV, ethanol, bleach, temperature), sterile filtered (0.22 μm filter), or autoclaved. The formulation was composed of 40×10^6^ C2C12 cells, 100 μL D-mannose (64.8% D-mannose in PBS), 80 μL DMEM, and 1740 μL ink per 2 mL. The bioprinted samples were incubated in 6-well plates with differentiation media. Cultured media was refreshed every 2-3 days. The width and length of samples were measured after printing and 1, 3, and 7 days in culture with a calliper.

### 2.13 Metabolic activity assay

Cell viability of the inks screening assays and the bioprinted samples was assessed using the 96®AQueous One Solution Cell Proliferation assay (MTS assay, Promega, USA). The supplier protocol was used. Briefly, the samples were rinsed and incubated for 3 h at 37 °C in a 5 % CO2 atmosphere with DMEM without phenol red and FBS and with HEPES, Glutamax and the MTS reagent (5:1 ratio). After incubation, 100 μL was transferred to a 96-well plate (triplicate) to measure their absorbance at 490 nm by a microplate reader (Synergy Biotek H1 Microtiter Plate Reader, Synergy Biotek). Each bioprinted sample was incubated with 2.5 ml of the reaction solution. For the study of Fig.2, three 15 μL samples were incubated with a volume of 0.5mL.

### 2.14 Cell morphology

After culture, the samples were rinsed thrice with sterile PBS and then fixed with formalin 4 % (v/v) formalin. The samples were rinsed with PBS, and the cells were permeabilized with 0.2 % Triton X-100 (Sigma-Aldrich, 93443) in 0.2 % PBS, with mild shaking for 15 min. Then, the samples were rinsed thrice with PBS and incubated with phalloidin–tetramethylrhodamine B isothiocyanate (Phalloidin-TRITC, 1:100, Invitrogen, D3571) for 30 min in the dark. The samples were rinsed and incubated with 4,6-diamino-2-phenyindole dilactate (DAPI, 1:1000, Molecular Probes, R415) for 15 minutes. The samples were observed by <u>confocal microscopy</u> (Zeiss LSM 780) and excited at 340 and 495 nm wavelengths. Emission was set at 570-620 nm, and the observation was carried out with 40 × and 63 x oil Zeiss objectives.

### 2.15 Immunocytochemistry

After 7 days in culture, the bioprinted samples were rinsed twice with sterile DPBS and fixed with 2.5 % formalin for 30 min at room temperature (24 °C). The samples were incubated for 15 min with 0.2 % Triton X-100. Then, the samples were washed thrice with PBS for 5 min and then incubated with 1 % bovine serum albumin (BSA), 22.52 mg/mL glycine in 0.1 % Tween 20 in PBS (PBST) for 45 min to block unspecific binding of the antibodies. The samples were incubated with mouse monoclonal to heavy chain Myosin/MYH3 (Abcam,ab264038, 1/500 in 1 % BSA) in PBST overnight at 4°C. The solution was aspired, and the samples were washed thrice with PBS for 5 min each. Finally, the samples were incubated with the goat polyclonal secondary antibody to Mouse IgG - Alexa Fluor® 555 (Abcam, ab150114, 1/500) in 1 % BSA solution for 2 hours at room temperature in the dark. The solution aspired, and the samples were washed three times with PBS for 5 min in the dark. The samples were observed by <u>confocal microscopy</u> (Zeiss LSM 780) and excited at the wavelengths of 555 nm. Emission was set at 565 nm, and the observation was carried out with 40 × and 63x oil Zeiss objectives.

### 2.16 Texture Profile Analysis

The hardness, cohesiveness, chewiness, and resilience of the acellular and cellular samples were analysed by Texture Profile Analysis (TPA) using a Shimadzu AGX-10 kN Texture Analyzer (Shimadzu, Japan) with a 50 N load cell and a cylindrical probe with a diameter of 50 mm. The experimental parameters included compression at a speed of 0.5 mm/s (equivalent to the rate of probe pulling) under a maximum strain of 50%.

### 2.17 Statistical Analysis

The DoE and associated statistical analyses were performed using TIBCO Statistica (version 10). All the other statistical analysis were performed using student’s *t*-test (unequal variance and one-tailed). The obtained p-values can be consulted in the supporting information.

## 3 Results

### 3.1 Screening shape fidelity of biopolymers and microstructure of the selected inks

Phase-separated gels can display flow behaviour and gel properties that can benefit bioprinting (Çak & Foegeding, 2011; Oliveira et al., 2020). This previous structuration approach was employed to screen different combinations of proteins and naturally derived polysaccharides. The qualitative printability of pea protein isolate (PPI), whey protein isolate (WPI – as control), brown rice protein (RP), soy protein isolate (SPI) and the polysaccharides gellan gum (GG), τ-carrageenan (τCar), κ-carrageenan (κCar) was assessed – Fig. S1, Fig. S2. The most promising combinations identified, based on the capacity to build up the layers, smoothness and absence of syneresis, were WPI-GG, WPI-τCar, RP-GG, RP-τCar, SPI-τCar and PPI-τCar. PPI-τCar was selected for further studies, considering pea protein’s inherent presence of phenolic compounds with antioxidant activity, which may benefit cultured cells (Faber et al., 2024).

One other strategy to mitigate the negative impacts of excessive shear stress on cells during printing is to use tapered nozzles instead of blunt end and long ones (Gunther et al., 2022). Considering this, tapered tips were used in the starting tests, and a custom nozzle for a droplet-wise introduction of ions during printing was fabricated – Fig. S3a. A commercial blend of calcium gluconate and calcium lactate (Gluco) was chosen as the source of calcium ions to promote *in situ* ionic self-assembling of ℩Car. This source is tasteless and has demonstrated slower gelation kinetics than pure calcium chloride. Compared with calcium chloride, the gelation time is 5 to 20 times slower in alginate^19^, permitting better membrane thickness and hardness control. Additionally, those calcium ions can be a more bioavailable source of calcium (Krause et al., 2015).

A factorial design experiment was conducted to understand the combinatorial effect of the PPI and τCar on the shape fidelity of the printed structures with 100 % infill – Table S1. The obtained model representing the relation of shape fidelity and PPI and τCar was:

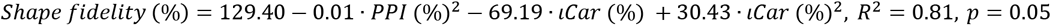

Fig. 1b presents the surface response, depicting the function, along with the top-view photographs of the printed samples and the water-sheared ones, confirming the suitability of the approach for producing independent fibres. The ability to create soft fibres was also confirmed with 10% SPP and 1% τcar ink, which could be gently separated with a spatula – Fig. S3b. In the range analyzed, the shape fidelity of the PPI-based inks was above 88 %. Increasing τCar concentration enhanced fidelity, regardless of PPI concentration. However, the same trend was not observed with PPI, which had a negative effect factor (−0.01).

TThe inks were stained with rhodamine B, which binds to the protein phase, and the microstructure was observed under confocal microscopy – Fig. 1c. Visual inspection revealed no major differences in the pea protein particle shape, which was irregular and three-dimensional with variable sizes. However, particle aggregation was observed in some areas, especially in the cases of fidelity below 90 %. The particles ranged from less than 1μm to up to 50μm and some particles of approximately 100μm were seen. The samples and controls selected for further studies were: 7 % PPI-1 % τCar (7P1C), 4 % PPI-1 % τCar (4P1C), 7 % PPI-1.5 % τCar (4P1.5C), 0 % PPI-1 % τCar (0P1C), 0 % PPI-1.5 % τCar (0P1.5C).

### 3.2 Rheological and fludic properties of PPI-τCar inks

Rheological tests were performed to assess the shear-thinning capacity, thixotropy, and stress levels upon printing – Fig. 1d,e and Fig. S4. In both flow curves and the thixotropy test, a synergistic effect of the combination of PPI and τCar was evident. The phase separation significantly increased the viscosity and the capacity to recover after high shear conditions. At 1000 s^-1^, the viscosity of 7P1C reached 205 mPa.s, and 1.5 s later, it recovered to 21 Pa.s. The initial viscosity was recovered in 56 s. The flow curve data was fitted with the Herschel-Bulkley model. The obtained fluidic properties are detailed in Table S2. The flow index of 7CP1C ink was 0.17, indicating a highly shear thinning behaviour. Similar behaviour (n<0.2) was observed for 4P1C, 4P1.5C, 1P1.5C, 0P1.5C, and 7P1C but not for 0P1C and 7P0C, which exhibited flow indices of 0.44 and 0.84, respectively. The consistency index, *K*, which correlates with the viscosity, did not exceed 35 Pa.s^n^. The lowest values were observed in the inks containing only ℩Car or 7PPI, supporting the previously mentioned synergy. The yield stress, τ_o_, which correlates with the force required to initiate ink flow, was negative in most of the cases. This negativity could be attributed to the high thixotropic behaviour introduced by ℩Car into the systems. In our experiments, the inks were subjected to pre-shear, such as in a real 3D printing scenario. Nevertheless, a small portion of the viscosity could still be under recovery, justifying the negative values.

### 3.3 Micro-extrusion and flow regimes limits of the inks and shear stress

Shear stress can affect cell viability, induce cell elongation, influence other intracellular mechanisms, or even lead to cell death if it exceeds certain thresholds (Boularaoui et al., 2020). The levels of shear stress are related to various factors, including nozzle geometry, printing speed and the viscoelastic properties of the ink. Nonetheless, the nozzle wall is one of the regions with the highest stress levels in the printing process (Blaeser et al., 2016; Boularaoui et al., 2020). The shear stress at the wall of the inks at different printing speeds was calculated using the equation described in the rheology methods – Fig. 1f.

For printing speeds up to 200 m/s, the shear stress remained below 92 Pa given a nozzle diameter of 1.07 mm (diameter used in further bioprinting tests). These values were significantly lower than the limits reported in the literature to avoid major cell death (Reina-Romo et al., 2021), which should preferably be below 300 Pa. The diameter of the nozzle substantially affects the results. Nonetheless, for a 200 μm nozzle, we estimate a stress of 245 Pa at 200 mm/s for the 7P1C ink.

To analyse the range of actual pressure required to print, the micro-extruded mass of the inks was measured using a pressure-controlled pneumatic system. The mass was converted to the theoretical printing speed and plotted against the applied pressure – Fig. 1g. At very low pressures 0.1 bar, the 7P1C flowed at a speed of 32.2±7.4 mm/s, while at 0.15 bar, it flowed at 160.1±2.4 mm/s. With evident turbidity, the printing speeds could reach over 1000 mm/s at high pressures. Thus, results confirm that very low pressures permited high flows, which can indicate low shear stress upon bioprinting.

Printability is often assessed in terms of shape fidelity or rheological properties. Besides shear stress limits, other underlying limiting physical events (Zhang et al., 2018) influenced by the viscoelastic properties, such as the actual flow regimes, should be considered. In particular, the Reynolds number, *Re*, is a dimensionless number used to determine if the fluid flow is laminar, transient or turbulent. The *Re* number was calculated for printing speeds up to 200 mm/s for the 7P1C ink and plotted against the printing speed and the corresponding extruded mass, delineating the thresholds of laminar, transient and turbulent flow - Fig. 1h. These findings suggested that 7P1C could be micro-extruded up to 120 mm/s in the laminar flow regime, with shear stress at the wall of 82 Pa, a shear rate of 902 s^-1^ and an apparent viscosity of 90 mPa.s.

### 3.4 Self-assembling and preliminary C2C12 myoblasts cytocompatibility assessment

The inks exhibited excellent flowability and maintained their shape integrity with a 100 % infill, as observed in the section 3.1. However, the stability was time-limited in phosphate-buffered saline solution (PBS), and delamination occurred – Fig. 2.

**Fig. 2.**
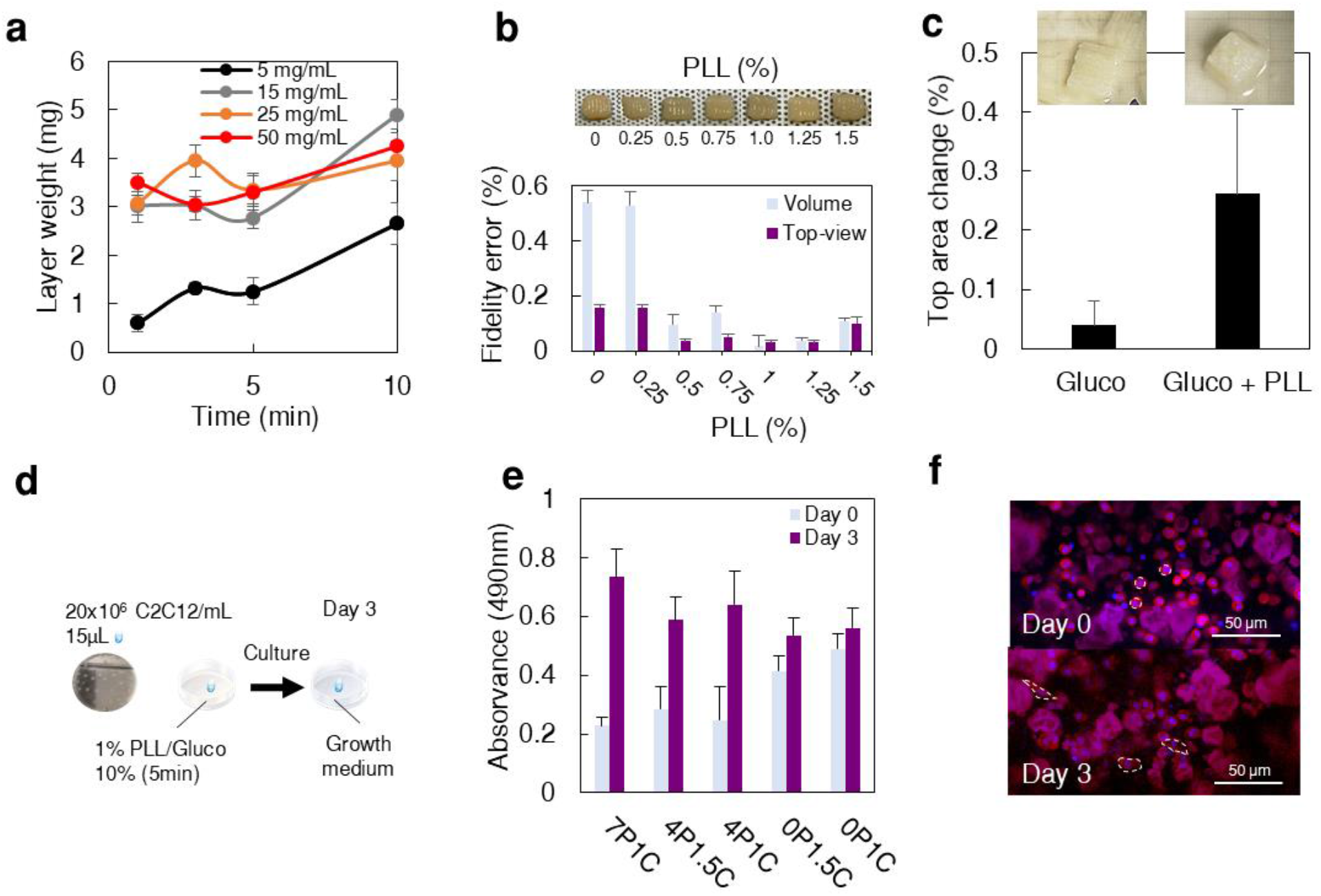
Preliminary assessment of C2C12 cytocompatibility in the inks with self-assembled PLL coating. (a) Effect of time and PLL concentration on the total mass deposited on 20μL droplets (n=4). (b) 7P1C samples printed at 10 mm/s structured by *in situ* ionic and polyelectrolyte self-assembling at various PLL concentrations, with 10 % Gluco and using a 1.07 mm blunt co-axial nozzle (n=3). Dimensional accuracy of the printed samples regarding volume and top-view area. (c) Top area change of the samples self-assembled only with Gluco (sample delamination) or Gluco and PLL after 6 days in culture media at 37 °C (n=3). (d) Schematic illustrating the preparation of the cellular droplets. (e) Metabolic activity was measured by MTS assay for C2C12 cells embedded in the inks after 0 and 3 days cultured in a growth medium (n=6). Three droplets were measured per well, equivalent to an initial cell concentration of 0.9×10^6^ cells per well. P-values of statistical analysis can be found in Table S3. (f) Confocal microscopy images of the morphology of C2C12 stained for cell nuclei (DAPI) and cytoplasm (TRITC-phalloidin), showing cell spreading after 3 days in culture. The data is represented as average and standard deviation.

The continuous phase of the inks, ιCar, carries two sulphate charges per monomer unit, potentially endowing the system with the polyelectrolyte complexation capacity. ε-Poly L-Lysine (PLL) was chosen as the positively charged polyelectrolyte to complex with ιCar aiming to increase shape stability. PLL is food-grade and has bacteriostatic properties. Previous reports have documented the complexation of PLL and ιCar (Wang et al., 2021).

The capacity of PLL, in the presence of 10 % Gluco, to coat the inks was evaluated, revealing an increase in the mass deposited on 15 μL drops over time and with increasing PLL concentration – Fig. 2a. Following this, the 7P1C samples were coaxially printed with different PLL concentrations, 0 to 1.5 %, along with 10 % Gluco, and the dimensional accuracy was quantified – Fig. 2b. While the shape fidelity improved with PLL concentration above 0.5 %, there was a noticeable increase in fibre dragging, suggesting diminished immediate interlayer adhesion, particularly with PLL concentrations above 1 %. Nonetheless, the combination of printing 7P1C with Gluco and PLL reduced delamination – Fig. 2c.

To our knowledge, the cytocompatibility of PPI and ιCar inks has not yet been reported. Therefore, preliminary *in vitro* studies were conducted using C2C12 murine myoblast cell line (Milasincic et al., 1996), which is an extensively used cell model to study myogenesis. C2C12 cells, at a concentration of 20×10^6^ cells/mL, were embedded in 7P1C and control inks forming 15 μL drops and cultured for up to 3 days – Fig. 2d. Metabolic activity and cell morphology were analyzed as depicted in Fig. 2d,e. Over the 3-day culture period, the metabolic activity of C2C12 cells increased, with more pronounced effects observed in the 7P1C samples. The metabolic activity of C2C12 cells was generally significantly higher on 7P1C ink than on the other samples after 3 days, except compared to 4P1C (p-value = 0.054). Initially, round cell morphologies transitioned to elongated and spread cytoplasms, indicating cellular mobility within the phase-separated inks. As anchorage-dependent cells, the results suggest cell attachment has occurred. Those observations also suggested that increasing PPI content was beneficial.

Factors potentially contributing to enhance cell activity include increased permeability, cell adhesion, mobility and spreading. Those could have been influenced by the ink’s mechanical and biochemical properties, the presence of the physical anchorage points and antioxidants that the structures of PPI could provide. Additionally, the sulphate groups from ιCar may have facilitated cell adhesion or sequestered adhesion factors from cultured media, thereby enhancing cell adhesion (Kowalczyńska & Nowak-Wyrzykowska, 2003). Therefore, we have considered that the ink could have the potential for bioprinting.

### 3.5 Ionic and polyelectrolyte in situ self-assembling bioprinting

A key constraint in developing thick cell-cultured structures is the permeability and diffusion of nutrients and cell metabolites. Limited diffusion is a cell stress source due to nutritional imbalance, low oxygen delivery, and poor metabolite removal (McMurtrey, 2016). Estimates suggest that the maximum hydrogel thickness for viable embedded cells can range between 400 to 1400 μm for glucose concentrations of 10-20 mM glucose (Tomiyama et al., 2020). The actual thickness will depend on diffusion dynamics, the structure of the gel and the specific requirements of the cells. Ensuring timely and proper cell nutrition is fundamental to optimizing cell activity, including their capacity for protein production.

To increase the permeability of the 3D structure, we designed a challenging single-wall prism with a porous structure, as illustrated in **Fig. 3**. As expected, the 7P1C shape fidelity was inferior without *in situ* crosslinking, as demonstrated in method 1. When self-assembled with 0.5 % PLL and 10 % Gluco, the 3D structure was unstable, and the wall started collapsing from the outside-in after 5-6 layers – method 2. Thus, the *in situ* ionic self-assembling of Ca2^+^ ions and the self-assembling of the PLL was insufficient to counteract the gravitational forces and the overlying weight.

**Fig. 3.**
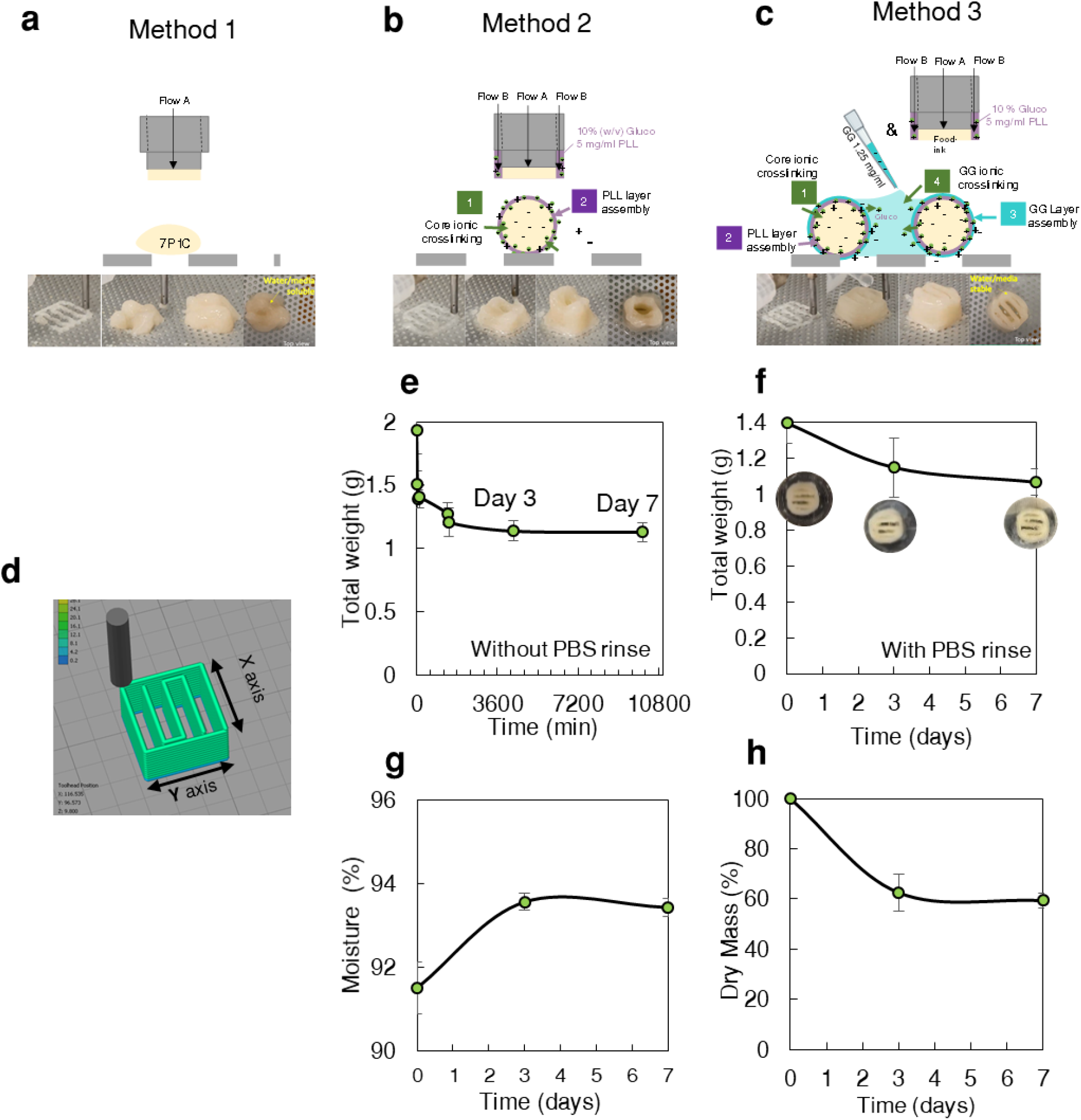
Schematic representation of the structuration and self-assembling methods and stability assessment of the phase-separated 7P1C ink co-axially printed at 10 mm/s. (a) Co-axial microextrusion-based printing of 7P1C. (b) Co-axial microextrusion printing by *in situ* self-assembling enabling PLL coating and ink gelation with calcium gluconate and calcium lactate with 10 % Gluco and 0.5 % PLL. (c) Coaxial microextrusion printing by *in situ* self-assembling enabling PLL coating and ink gelation, followed by self-assembling with GG and also forming a fluid gel in the open pores. (d) Sliced model model (x= 15.4, y=15.4 and z= 9.8 mm). (e) Total weight of the samples immediately post printing (without PBS rinse) and after incubation in PBS/0.2 % sodium azide for 30, 60 min and 1, 3 and 7 days at 37. (f) Total weight of the samples immediately post printing (with PBS rinse) and incubation in PBS/0.2 % sodium azide for up to 7 days at 37 °C. Top view of the samples after incubation. (g) The moisture content of the samples. (h) Total dried mass of the incubated and freeze-dried samples. Data is presented as average and standard deviation, n=5.

To overcome this, we applied a polyelectrolyte water-based solution over each printed layer to enhance interlayer binding and provide internal physical support to printed filaments. It contained 1.25 mg/mL of gellan gum (GG, pH 7), an extremely fluid gel, and a negatively charged polyelectrolyte (Milasincic et al., 1996). GG is a versatile food hydrocolloid known to form fluid gels, or structured liquids, at low concentrations such as 1.25 mg/mL (Milasincic et al., 1996). The gel strength was reported to be approximately 20 g when crosslinked with 0.6 % calcium and 100 g with 0.15 % calcium.

As a result, with method 3, a weak gel was formed inside the pores through the self-assembly with calcium ions from Gluco that the co-axial nozzle had previously deposited. The negatively charged GG could self-assemble with the positively charged PLL-surface coated filaments of the underlying layers. Given that the subjacent layers were deposited with a PLL coating, they could also self-assemble with the PLL-GG-coated subjacent layers, potentially enhancing 3D shape stability. And, indeed, a reduction in the fibre sagging was observed.

The stability of the 7PC1 samples (method 3) in PBS was analyzed. The weight of both rinsed and unrinsed samples and the total dried mass and moisture content up to day 7 are illustrated in **Fig. 3e,f**. It was observed that the total weight of both rinsed and unrinsed samples decreased over time, indicating these structures did not swell. The most significant decrease occurred within the initial hours, especially in the unrinsed samples, which is believed to be attributed to the release of the GG fluid gel from the 3D structure. After 3 days, the sample weight, moisture content and dry mass stabilized at approximately 1.06 g (rinsed) or 1.13g (unrinsed), with a moisture content of 93.4 % and a dried mass loss of 59.4 % - Fig. 3g,h. According to the build summary generated by simplify3D, the volume of the sliced model is 1.05 ml, theoretically corresponding to 84 mg of solids, excluding *in situ* crosslinking. After printing, the total solids were 128 ± 2mg, decreasing to 74 ± 8 mg after 3 days and 70 ± 3.4 mg after 7 days. This represents a 30 % variation compared to the theoretical model, which could be related to some ink trapped in the printing surface, small variations in the actual printed volume and solids loss during the stability tests.

### 3.6 Characterization of the 7-days differentiated bioprinted samples: morphology, texture and protein

The bioinks composed of samples with 7P1C and 20×10^6^ cells were bioprinted and cultured for 7 days in differentiation media – Fig. 4a. Preliminary tests were conducted to assess shape fidelity and validate the potential methodology – Fig. 4b. The dimensional accuracy was generally lower in the 7P1C bioprinted samples (20×10^6^ C2C12/mL) except for the Z axis – Table S4, Fig. 4b. The cells only reduced the sample length, and particularly on the X axis, by approximately 6 % compared to the acellular 3D samples. One possible explanation is that the longest fibres in the printing direction align predominantly along the X-axis, and cells induced shrinkage. Interestingly, the embedded cells increased the height accuracy by 5.4 %, permitting a height of 8.5 mm after 7 days in culture. This suggested cell assembling and interactions with ink, increasing 3D structure integrity.

**Fig. 4.**
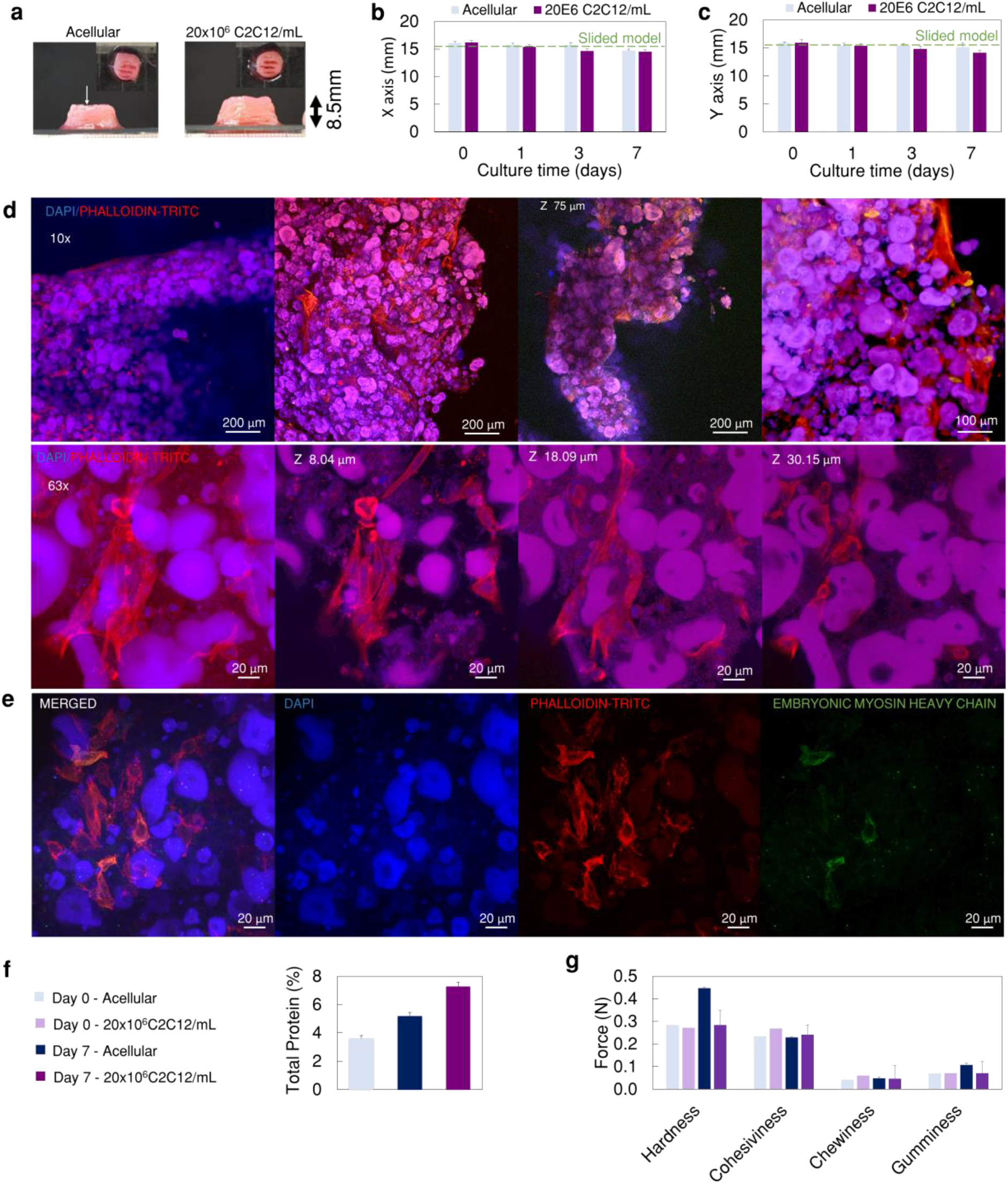
Characterization of the bioprinted samples cultured in differentiation medium for 7 days. (a) Photographies of the side and top-view of the acellular and cellular printed samples. (b, c) Length of the samples during the culture time on the X and Y axis (n=3). (d) Confocal microscopy images of the samples stained with phalloidin-TRITC (red, cytoskeleton) and DAPI (blue, nuclei) showing the cell networks. (e) Confocal immunocytochemistry images for embryonic MyHC (green), nucleus and cytoskeleton. (f) Total protein with and without cells, quantified by Kjeldahl analysis (conversion factor 6.25), (n=3). (g) Texture properties of the bioprinted samples (n=3). Data is presented as average and standard deviation.

Notably, we have not observed the typical high sample shrinkage (Ianovici et al., 2022), potentially due to the phase separation. We hypothesise that the cells did not fully occupy the continuous phase (℩Car) or the particles were hard enough to contrapose the cellular contraction forces, or a combination of both. Further studies are required to confirm whether the irregular 3D-shaped protein structures were indeed a key factor contributing to the shape stability and when typical cell culture anchorages (Kang et al., 2021) would be required.

The MTS assay revealed cells were metabolically active after 7 days of differentiation – Fig. S5. By confocal analysis, it was evident that there was extensive cell spreading and formation of cellular networks between the pea particles – Fig. 4d.

During differentiation, myoblasts become mononuclear myocytes, which fuse into myotubes. Several myosin-heavy chain (MyHC) isoforms are expressed during muscle tissue formation or myoblast differentiation (Brown et al., 2012). The embryonic MyHC, also known as myosin heavy chain 3 (emb-MyHC, encoded by Myh3 gene), is highly expressed in early myoblasts differentiation (Tang et al., 2023) but not in myoblasts (Brown et al., 2012) or adult muscle except during muscle regeneration or disease (Agarwal et al., 2020). Although relatively unexplored, it plays crucial roles in regulating muscle fibre size, number, and type, as well as myogenic progenitor and myoblast differentiation (Agarwal et al., 2020). The absence of emb-MyHC leads to the misregulation of muscle differentiation-related genes and several diseases. Notably, *in vitro* knock-down of the Myh3 gene accelerates the initial differentiation rate after 3-5 days, but the rate later declines. Patterns of embryonic/neonatal and adult MyHC isoforms have been identified *in vitro* in C2C12 cells (Brown et al., 2012), considering emb-MyHC an early isoform. Therefore, emb-MyHC was the target chosen to identify early differentiation. The immunocytochemistry performed on the bioprinted samples after 7 days in culture revealed the presence of emb-MyHC in the cytoplasm in several cells, validating the existence of cells in the early differentiation stage – Fig. 4e.

While the protein contribution from adult myofibers is reported as 675 pg per cell nuclei in rat leg muscle (Wiśniewski et al., 2014), mono-nucleated myoblasts are expected to have less protein. In fact, 20 to 30 % (200-300 pg) is the typical cell protein concentration reported for several mononucleated cells (Wiśniewski et al., 2014). Consequently, although the bioprinting was performed at a high cell concentration (20 million/mL), the protein amount introduced by the cells per sample can be estimated as just 4.2 mg for 200 pg/cell. Understanding how to improve cellular protein production through ink design, bioprinting control, and culture is fundamental.

According to Kjeldahl analysis performed by a certified laboratory, the initial protein content in the acellular samples was 3.62 % protein. After culture, it increased to 5.2 % in the acellular samples, suggesting protein adhesion to the PLL/GG coated surfaces or incomplete removal of the culture medium. Nonetheless, the protein content was significantly higher in the bioprinted samples, comprising 7.3 % protein. Subtracting the protein of the cellular and the acellular samples, the result is 2.1 g/100 g. Considering that 1) there was no significant pea protein loss during culture; and 2) the bioprinted sample volume after 7 days was 1.05 mL, then each bioprinted sample contained 76.4 mg of cell-produced protein. Each sample initially contained 21×10^6^ cells, meaning that each cell yielded 3638 pg of protein, representing an 18-fold protein increase per initial cell over 7 days.

Texture analysis, however, did not reveal an improvement with culture, and all the 3D samples were very soft. In particular, the hardness of the bioprinted samples was 283 mN, similar to the values observed before culture. The exception was a slight increase in the hardness of the acellular samples, possibly caused by the height reduction and the packing of the printed structure.

## 4 Conclusion

Here, we reported a proof-of-concept for a novel food-grade, animal-free ink and a bioprinting method integrating *in situ* ionic and polyelectrolyte self-assembly with fluid gels. This innovation enabled the structuration of low-shear stress and low-viscosity food-grade ingredients supporting myoblasts’ early differentiation and cellular protein production. After the differentiation period of 7 days, long multi-cellular structures (>200 μm) were observed, suggesting myoblast differentiation and fusion around the particles. Early differentiation was also confirmed by emb-MyHC in the cells’ cytoplasm and we did not observe sample shrinkage, potentially due to the phase separation.

The method successfully allowed us to created single-wall bioprinted constructs that were shape-stable during the culture period with the 7P1C ink. The resulting bioprinted samples exhibited a soft texture (hardness of 283 mN), which allowed us to isolate the effect of scaffolding on the texture. Nevertheless, we consider our study a proof-of-concept that will benefit from further in-depth studies.

While some studies have already explored bioprinting for cultivated fish/meat applications, summarised in Table 1, few have developed food-grade bioinks or studied their effect on texture and nutrition. To our knowledge, this study is the first to report a food-grade ink observing an increase in cellular-derived protein of up to 18-fold/per initial cell after 7 days of differentiation. Bioprinting was performed at the theoretical flow of 32.2 g/h, significantly above the current published research (0.56 to 18 g/h). Nevertheless, a C2C12 myoblasts cell line and non-animal-free cultured media were used. Future research should utilize a more product-representative cell line and animal-free culture media alternatives, e.g., replacing albumin and FBS.

Compared with FRESH-based approaches, which are the most frequently used to structure low-viscosity ink, our method may have specific limitations regarding resolution and a minimum printable viscosity that have yet to be determined. Future studies should also explore the optimization of printing speeds, with an interest in achieving a speed of up to 120 mm/s while maintaining the laminar flow without resorting to narrow nozzles, which could hinder product flow for food applications.

This integrative bioprinting by *in situ* self-assembling provides a novel approach to potentially produce robust, cell-dense platforms for further studies in cell-based foods. In addition, the effect of different variables (e.g., biopolymers ratio and type, particle size, nozzle, speed) can be studied to increase the knowledge on the effect of bioprinting, bioink, and culture on, e.g., the texture and nutrition of cell-based food.

## CRediT authorship contribution statement

**Sara M. Oliveira:** Conceptualization, Data curation, Formal analysis, Investigation, Methodology, Writing – original draft, review & editing, Supervision, Project administration. **Gabriel**

**DeSantis:** Investigation, Writing – review & editing. **Lorenzo M. Pastrana:** Supervision, Resources, Methodology, Writing – review & editing.

## Acknowledgements

This work was supported by The Good Food Institute through the project M3atD - 3D bioprinting for cultivated meat and the European Union through the project FEASTS - Fostering European Cellular Agriculture for Sustainable Transition Solutions (Grant agreement ID: 101136749). The authors also acknowledge the Fulbright Commission Portugal for the Fulbright/FLAD Open Study/Research Award granted to Gabriel DeSantis. We also would like to thank Pedro M. Silva for helping with cell harvesting for the bioprinting studies. We also would like to thank the Nanophotonics and Bioimaging (NBI) Facility from INL for their support, especially to Mariana T. Carvalho.

## Appendix A. Supplementary data

The following is the Supplementary data to this article.

## Data availability

Data will be made available on request.

## References

Agarwal, M., Sharma, A., Kumar, P., Kumar, A., Bharadwaj, A., Saini, M., Kardon, G., & Mathew, S. J. (2020). Myosin heavy chain-embryonic regulates skeletal muscle differentiation during mammalian development. Development (Cambridge), 147(7), 1–14. 10.1242/dev.184507

Blaeser, A., Duarte Campos, D. F., Puster, U., Richtering, W., Stevens, M. M., & Fischer, H. (2016). Controlling Shear Stress in 3D Bioprinting is a Key Factor to Balance Printing Resolution and Stem Cell Integrity. Advanced Healthcare Materials, 5(3), 326–333. 10.1002/adhm.201500677

Boularaoui, S., Al Hussein, G., Khan, K. A., Christoforou, N., & Stefanini, C. (2020). An overview of extrusion-based bioprinting with a focus on induced shear stress and its effect on cell viability. Bioprinting, 20(August), e00093. 10.1016/j.bprint.2020.e00093

Brown, D. M., Parr, T., & Brameld, J. M. (2012). Myosin heavy chain mRNA isoforms are expressed in two distinct cohorts during C2C12 myogenesis. Journal of Muscle Research and Cell Motility, 32(6), 383–390. 10.1007/s10974-011-9267-4

Çak, E., & Foegeding, E. A. (2011). Combining protein micro-phase separation and protein e polysaccharide segregative phase separation to produce gel structures. Food Hydrocolloids, 25, 1538–1546. 10.1016/j.foodhyd.2011.02.002

Dutta, S. D., Ganguly, K., Jeong, M. S., Patel, D. K., Patil, T. V., Cho, S. J., & Lim, K. T. (2022). Bioengineered Lab-Grown Meat-like Constructs through 3D Bioprinting of Antioxidative Protein Hydrolysates. ACS Applied Materials and Interfaces, 14(30), 34513–34526. 10.1021/acsami.2c10620

Faber, I., Pouvreau, L., Jan, A. Goot, V. Der, & Keppler, J. (2024). Modulating commercial pea protein gel properties through the addition of phenolic compounds. Food Hydrocolloids, 154(November 2023), 110123. 10.1016/j.foodhyd.2024.110123

Gunther, R. T., Seppala, J. E., Gunther, R. T., & Seppala, J. E. (2022). Simulated stress mitigation strategies inembedded bioprinting. Physics of Fluids, 34, 083112. 10.1063/5.0102573

Guo, X., Wang, D., He, B., Hu, L., & Jiang, G. (2023). 3D Bioprinting of Cultured Meat: A Promising Avenue of Meat Production. Food and Bioprocess Technology. 10.1007/s11947-023-03195-x

Hinton, T. J., Jallerat, Q., Palchesko, R. N., Park, J. H., Grodzicki, M. S., Shue, H. J., Ramadan, M. H., Hudson, A. R., & Feinberg, A. W. (2015). Three-dimensional printing of complex biological structures by freeform reversible embedding of suspended hydrogels. Science Advances, 1(9), e1500758. 10.1126/sciadv.1500758

Ianovici, I., Zagury, Y., Redenski, I., Lavon, N., & Levenberg, S. (2022). 3D-printable plant protein-enriched scaffolds for cultivated meat development. Biomaterials, 284, 121487.

Jeong, D., Seo, J. W., Lee, H., Jung, W. K., & Park, Y. H. (2022). Efficient Myogenic / Adipogenic Transdifferentiation of Bovine Fibroblasts in a 3D Bioprinting System for Steak-Type Cultured Meat Production. Advanced Science, 2202877, 1–16. 10.1002/advs.202202877

Kang, D.-H., Louis, F., Liu, H., Shimoda, H., Nishiyama, Y., Nozawa, H., Kakitani, M., Takagi, D., Kasa, D., Nagamori, E., Irie, S., Kitano, S., & Matsusaki, M. (2021). Engineered whole cut meat-like tissue by the assembly of cell fibers using tendon-gel integrated bioprinting. Nature Communications, 12(1), 5059. 10.1038/s41467-021-25236-9

Kowalczyńska, H. M., & Nowak-Wyrzykowska, M. (2003). Modulation of adhesion, spreading and cytoskeleton organization of 3T3 fibroblasts by sulfonic groups present on polymer surfaces. Cell Biology International, 27(2), 101–114. 10.1016/S1065-6995(02)00290-1

Krause, M., Keller, J., Beil, B., Driel, I. Van, Zustin, J., & Register, B. (2015). Calcium gluconate supplementation is effective to balance calcium homeostasis in patients with gastrectomy. Osteoporos Int, 26, 987–995. 10.1007/s00198-014-2965-1

Lawrie, R. A., & Ledward, D. (2014). Lawrie’s meat science. Woodhead Publishing.

Li, Y., Liu, W., Li, S., Zhang, M., Yang, F., & Wang, S. (2021). Porcine skeletal muscle tissue fabrication for cultured meat production using three-dimensional bioprinting technology. Journal of Future Foods, 1(1), 88–97. 10.1016/j.jfutfo.2021.09.005

Liu, W., Heinrich, M. A., Zhou, Y., Akpek, A., Hu, N., Liu, X., Guan, X., Zhong, Z., Jin, X., Khademhosseini, A., & others. (2017). Extrusion bioprinting of shear-thinning gelatin methacryloyl bioinks. Advanced Healthcare Materials, 6(12), 1601451.

Ma, Y., & Zhang, L. (2022). Formulated food inks for extrusion-based 3D printing of personalized foods: a mini review. Current Opinion in Food Science, 44, 100803. 10.1016/j.cofs.2021.12.012

Marques, D. M. C., Silva, J. C., Serro, A. P., Cabral, J. M. S., Sanjuan-Alberte, P., & Ferreira, F. C. (2022). 3D Bioprinting of Novel κ-Carrageenan Bioinks: An Algae-Derived Polysaccharide. Bioengineering, 9(3). 10.3390/bioengineering9030109

McMurtrey, R. J. (2016). Analytic models of oxygen and nutrient diffusion, metabolism dynamics, and architecture optimization in three-dimensional tissue constructs with applications and insights in cerebral organoids. Tissue Engineering - Part C: Methods, 22(3), 221–249. 10.1089/ten.tec.2015.0375

Milasincic, D. J., Dhawan, J., & Farmer, S. R. (1996). Anchorage-dependent control of muscle-specific gene expression in C 2 C 12 mouse myoblasts. In Vitro Cellular & Developmental Biology-Animal, 32, 90–99.

Mirdamadi, E., Tashman, J. W., Shiwarski, D. J., Palchesko, R. N., & Feinberg, A. W. (2020). FRESH 3D Bioprinting a Full-Size Model of the Human Heart. ACS Biomaterials Science and Engineering, 6(11), 6453–6459. 10.1021/acsbiomaterials.0c01133

Oliveira, S. M., Fasolin, L. H., Vicente, A. A., Fuciños, P., & Pastrana, L. M. (2020). Printability, microstructure, and flow dynamics of phase-separated edible 3D inks. Food Hydrocolloids, 109(June). 10.1016/j.foodhyd.2020.106120

Oliveira, S. M., Reis, R. L., & Mano, J. F. (2015). Towards the design of 3D multiscale instructive tissue engineering constructs: Current approaches and trends. Biotechnology Advances, 33(6). 10.1016/j.biotechadv.2015.05.007

Reina-Romo, E., Mandal, S., Amorim, P., Bloemen, V., Ferraris, E., & Geris, L. (2021). Towards the Experimentally-Informed In Silico Nozzle Design Optimization for Extrusion-Based Bioprinting of Shear-Thinning Hydrogels. Frontiers in Bioengineering and Biotechnology, 9(August), 1–14. 10.3389/fbioe.2021.701778

Song, W.-J., Liu, P.-P., Meng, Z.-Q., Zheng, Y.-Y., Zhou, G.-H., Li, H.-X., & Ding, S.-J. (2022). Identification of porcine adipose progenitor cells by fluorescence-activated cell sorting for the preparation of cultured fat by 3D bioprinting. Food Research International, 162, 111952. 10.1016/j.foodres.2022.111952

Tang, J., Yang, B., Song, G., Zhang, X., Wang, Z., Mo, Z., Zan, L., & Wang, H. (2023). Effect of bovine myosin heavy chain 3 on proliferation and differentiation of myoblast. Animal Biotechnology, 34(9), 4337–4346. 10.1080/10495398.2022.2149549

Tomiyama, A. J., Kawecki, N. S., Rosenfeld, D. L., Jay, J. A., Rajagopal, D., & Rowat, A. C. (2020). Bridging the gap between the science of cultured meat and public perceptions. Trends in Food Science and Technology, 104, 144–152. 10.1016/j.tifs.2020.07.019

Voutila, L. (2009). Properties of intramuscular connective tissue in pork and poultry with reference to weakening of structure. 94.

Wang, L., Zhang, C., Zhang, J., Rao, Z., Xu, X., Mao, Z., & Chen, X. (2021). Epsilon-poly-L-lysine: Recent Advances in Biomanufacturing and Applications. Frontiers in Bioengineering and Biotechnology, 9(September), 1–21. 10.3389/fbioe.2021.748976

Wiśniewski, J. R., Hein, M. Y., Cox, J., & Mann, M. (2014). A “proteomic ruler” for protein copy number and concentration estimation without spike-in standards. Molecular and Cellular Proteomics, 13(12), 3497–3506. 10.1074/mcp.M113.037309

Xu, E., Niu, R., Lao, J., Zhang, S., Li, J., Zhu, Y., Shi, H., Zhu, Q., & Chen, Y. (2023). Tissue-like cultured fish fillets through a synthetic food pipeline. NPJ Science of Food, 7(17). 10.1038/s41538-023-00194-2

Zhang, Z., Jin, Y., Yin, J., Xu, C., Xiong, R., Christensen, K., Ringeisen, B. R., Chrisey, D. B., & Huang, Y. (2018). Evaluation of bioink printability for bioprinting applications. Applied Physics Reviews, 5(4). 10.1063/1.5053979

Zhao, Y., Li, Y., Mao, S., Sun, W., & Yao, R. (2015). The influence of printing parameters on cell survival rate and printability in microextrusion-based 3D cell printing technology. Biofabrication, 7(4). 10.1088/1758-5090/7/4/045002

