## Supporting Information for "3D bioprinting of low-viscosity phase-separated food-grade bioinks by *in situ* self-assembly"

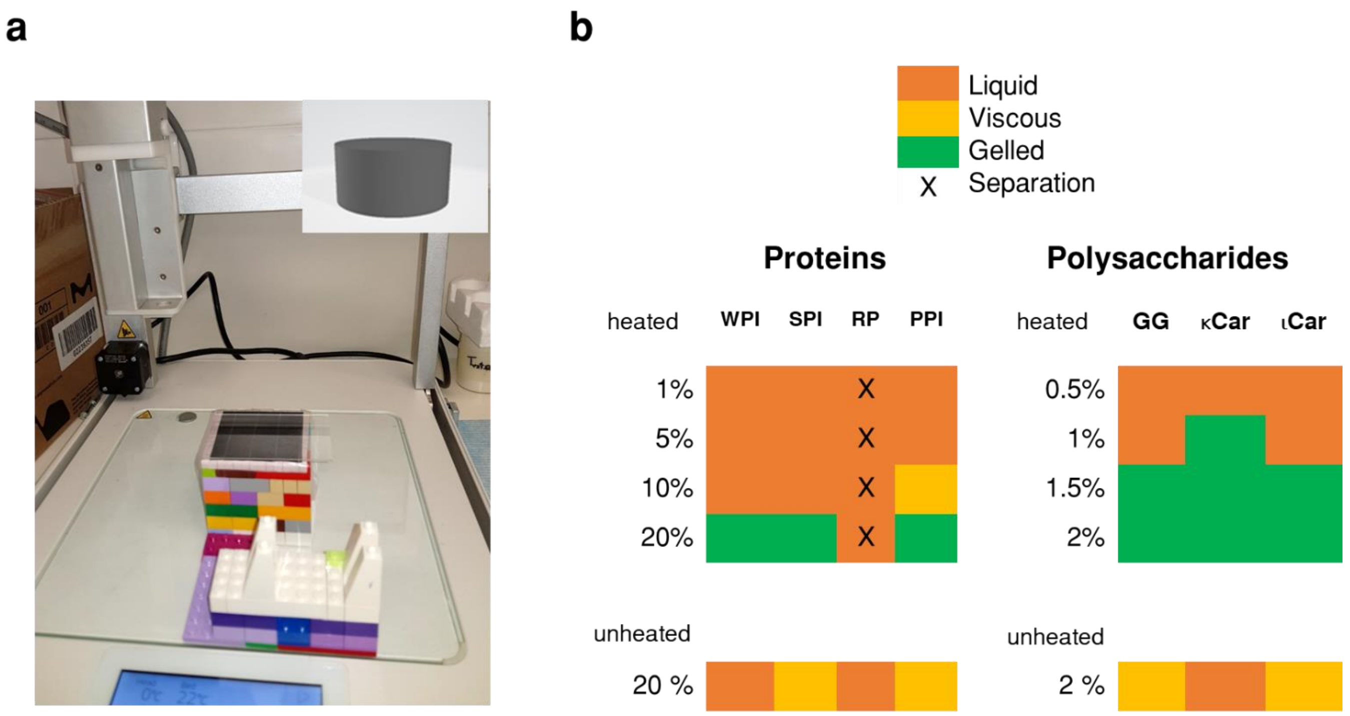

**Fig. S1**. (a) Printing setup for initial screening of different proteins and polysaccharides. (b) Qualitative evaluation of the printability of different proteins and polysaccharides.

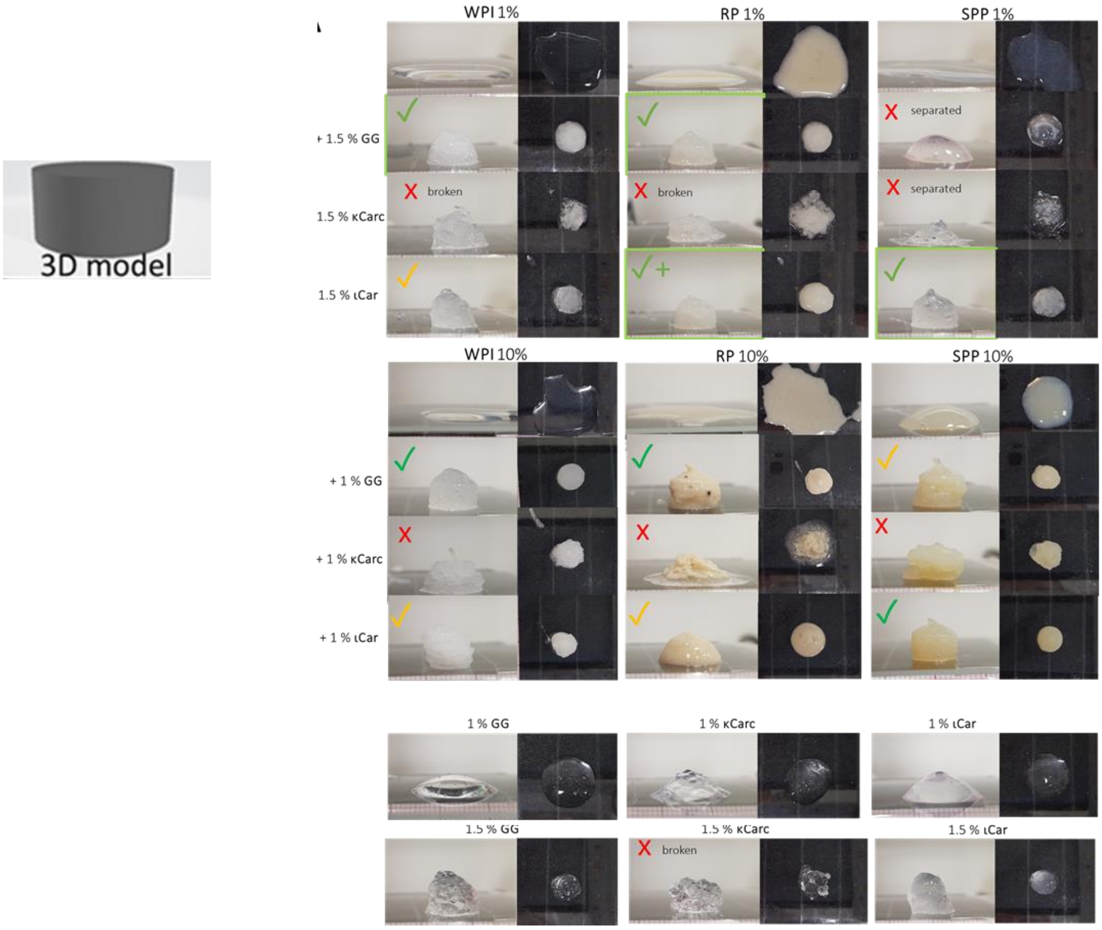

**Fig. S2.** Screening printability tests of the different proteins and polysaccharides. Without polysaccharides, all the proteins were in liquid form and not printable, independently of the concentrations tested. Some conditions presented complexation or the formation of strong and brittle gels of poor printability (e.g., 1% SPI- 1.5% κCar) or extensive syneresis (e.g., 10% RP-1% κCar). PPI displayed similar results compared with SPI at high concentrations.

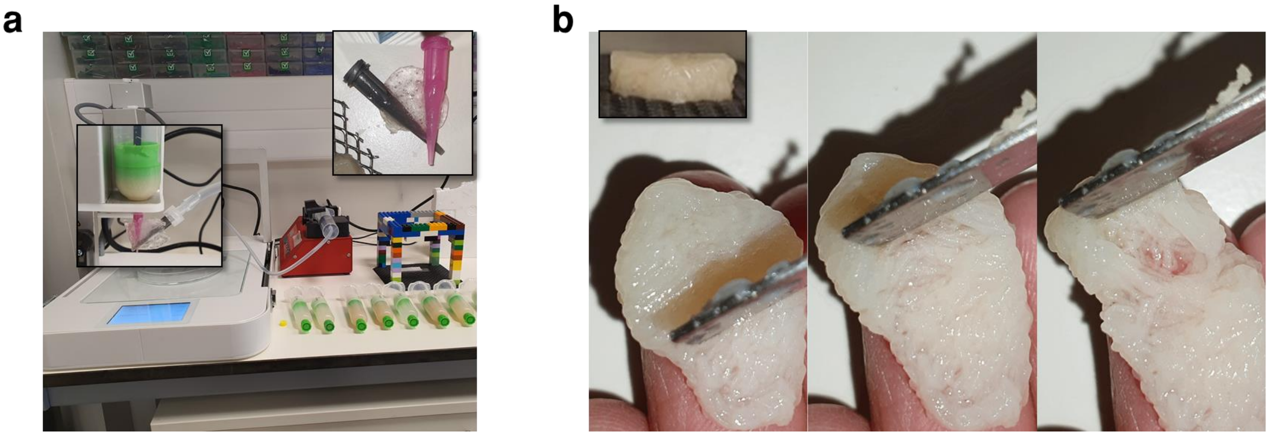

**Fig. S3**. Printing soft inks with a droplet-base system. (a) The printing setup features a custom-made nozzle with tapered tips for *in situ* crosslinking with calcium gluconate and calcium lactate (Gluco). (b) Soft fibre structures printed from 10% SPP ιCar ink printed at 30 mm/s using the exact structuration mechanism, revealed after opening with a spatula.

**Table S1.** Formulations tested for the central composite design, the natural values of the two independent variables studied were PPI (%) and ιCar (**%**). Output responses were height, width, and length, which were used to calculate printability. Output data is presented as the average of three replicates and the respective standard deviation (SD).

| PPI (%) | ιCar (%) | Height | SD | Width | SD | Length | SD | Printability |
|---|---|---|---|---|---|---|---|---|
| **1.5** | **1** | 3.9583 | 0.2305 | 18.42 | 1.07 | 22.62 | 0.29 | 91.8 |
| **1.5** | **1.5** | 4.6433 | 0.2964 | 19.095 | 0.305 | 19.21 | 0.22 | 92.7 |
| **8** | **1** | 4.0567 | 0.2148 | 17.32 | 0.25 | 17.195 | 0.135 | 90.5 |
| **8** | **1.5** | 4.2083 | 0.1087 | 18.855 | 0.265 | 18.355 | 0.355 | 94.5 |
| **0.15** | **1.25** | 4.4283 | 0.7274 | 18.855 | 0.035 | 19.405 | 0.375 | 94.4 |
| **9.35** | **1.25** | 4.2067 | 0.2926 | 17.62 | 0.46 | 17.16 | 0.43 | 90.43 |
| **4.75** | **0.9** | 3.8267 | 0.492 | 17.41 | 1.34 | 18.305 | 0.065 | 90.6 |
| **4.75** | **1.6** | 4.0983 | 0.4234 | 19.175 | 0.225 | 18.97 | 0.27 | 96.9 |
| **4.75** | **1.25** | 4.495 | 0.298 | 16.795 | 0.525 | 17.98 | 0.11 | 88.1 |
| **4.75** | **1.25** | 4.59 | 0.3681 | 17.38 | 1.79 | 18.08 | 0.48 | 88.4 |
| **4.75** | **1.25** | 4.4483 | 0.247 | 17.275 | 0.055 | 18.01 | 0.41 | 89.3 |

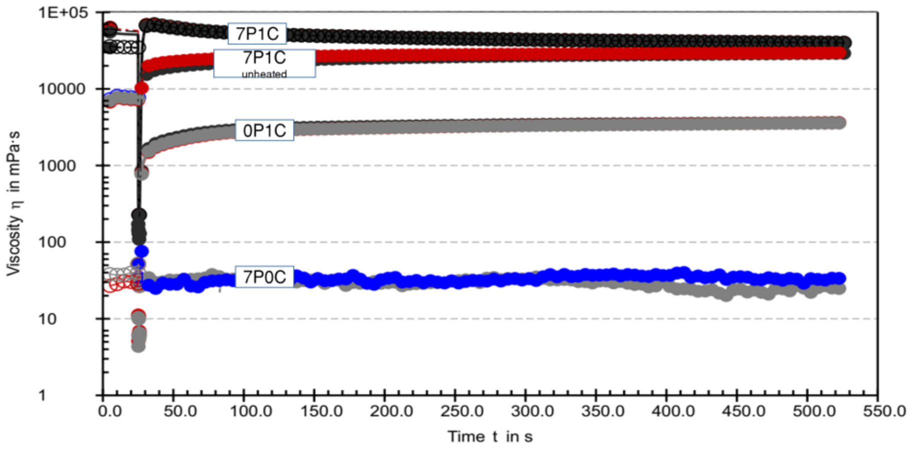

**Fig. S4.** Three-point thixotropy test of the 7P1C heated and non-heated inks and controls 0P1C and 7P0C (n=3).

**Table S2.** Fluidic parameters of the Herschel-Bulkley obtained for the PPI-ιCar inks. Data is presented as average and standard deviation (n=3).

| PPI (%) | ιCar (%) | $\tau_0$ (Pa) | SD | K (Pa.$s^n$) | SD | n | SD |
|---|---|---|---|---|---|---|---|
| **4** | **1** | -9.56329 | 0.31509 | 13.34689 | 0.175676 | 0.193701 | 0.0035 |
| **4** | **1.5** | -11.1641 | 1.325879 | 26.97118 | 1.589999 | 0.185292 | 0.00561 |
| **1** | **1.5** | -22.8001 | 0.744727 | 34.18181 | 0.494651 | 0.127682 | 0.00408 |
| **0** | **1** | -0.34795 | 0.005776 | 0.544463 | 0.013982 | 0.445892 | 0.003085 |
| **0** | **1.5** | -11.1495 | 0.833293 | 21.30583 | 0.715788 | 0.148454 | 0.003478 |
| **7** | **1** | -19.8949 | 1.471553 | 31.31811 | 1.596379 | 0.172662 | 0.00388 |
| **7** | **0** | 0.003144 | 0.001153 | 0.019891 | 0.001449 | 0.847616 | 0.002562 |

**Table S3.** Statistical p-values resulting from the single-tailed t-test considering unequal variance from the data of Fig. 2e.

| p-values | | | | | | |
|---|---|---|---|---|---|---|
| | | Day 3 | | | | |
| | | 7P1C | 4P1.5C | 4P1C | 0P1.5C | 0P1C |
| **Day 1** | 7P1C | 1.93635E-07 | 3.95752E-06 | 8.72089E-06 | 8.48614E-07 | 2.0111E-06 |
| | 4P1.5C | 9.50366E-08 | 6.52369E-06 | 9.41254E-06 | 1.62962E-05 | 9.09147E-06 |
| | 4P1C | 2.57973E-07 | 1.29444E-05 | 7.09881E-06 | 5.40195E-05 | 2.54476E-05 |
| | 0P1.5C | 2.96159E-06 | 0.000368102 | 0.000421394 | 0.000874697 | 0.000511128 |
| | 0P1C | 3.5421E-05 | 0.010009843 | 0.005048606 | 0.072346453 | 0.024465918 |
| | | Day 1 | | | | |
| | | 7P1C | 4P1.5C | 4P1C | 0P1.5C | 0P1C |
| **Day 1** | 7P1C | x | 0.063298372 | 0.356087074 | 2.77756E-06 | 5.24737E-09 |
| | 4P1.5C | x | x | 0.241578617 | 0.001985695 | 8.02479E-05 |
| | 4P1C | x | x | x | 0.002579867 | 0.000235499 |
| | 0P1.5C | x | x | | x | x |
| | 0P1C | x | x | x | x | x |
| | | Day 3 | | | | |
| | | 7P1C | 4P1.5C | 4P1C | 0P1.5C | 0P1C |
| **Day 3** | 7P1C | x | 0.003659012 | 0.054078236 | 0.000200431 | 0.00079513 |
| | 4P1.5C | x | x | 0.175795148 | 0.097721173 | 0.258032088 |
| | 4P1C | x | x | x | 0.028215781 | 0.073932871 |
| | 0P1.5C | x | x | x | x | 0.241663155 |
| | 0P1C | x | x | x | x | x |

**Table S4.** Dimensional accuracy of cultured samples. The dimensional accuracy was calculated as the percentage variation relative to the slide model. The p-value indicated compares the acellular and cellular samples for each axis and was calculated using a single-tailed t-test considering unequal variance. Data are presented as average and standard deviation (n=3).

| ***Sample*** | *Dimensional accuracy per axis (%)* | | |
|---|---|---|---|
| | **X** | **Y** | **Z** |
| **Cellular - 20x10$^6$ C2C12/mL** | 90.8±2.7 | 90.4±2.5 | 85.2±2.6 |
| **Acellular** | 96.3±3.1 | 92.3±4.4 | 79.8±2.7 |
| ***p-value*** | 0.067 | 0.314 | 0.053 |

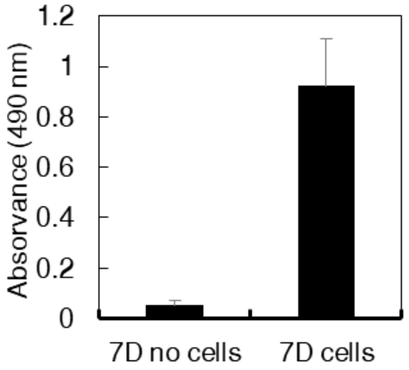

**Fig. S5.** Metabolic activity of acellular and cellular bioprinted samples after 7 days in differentiation media, measured by MTS assay (n=3). The data presented is the average and standard deviation.
